# Proteome partitioning constraints on long-term laboratory evolution

**DOI:** 10.1101/2023.09.08.556843

**Authors:** Matteo Mori, Vadim Patsalo, James R. Williamson, Matthew Scott

## Abstract

Adaptive laboratory evolution experiments provide a controlled context in which the dynamics of selection and adaptation can be followed in real-time at the single-nucleotide level^1^. And yet this precision introduces hundreds of degrees-of-freedom as genetic changes accrue in parallel lineages over generations^2^. On short timescales, physiological constraints have been leveraged to provide a coarse-grained view of bacterial gene expression characterized by a small set of phenomenological parameters^3–5^. Here, we ask whether this same framework, operating at a level between genotype and fitness, informs physiological changes that occur on evolutionary timescales. Using Lenski’s Ara-1 lineage adapted to growth in glucose minimal medium^6^, we find that the proteome is substantially remodeled over 40 000 generations. We apply our existing quantitative proteomics analysis to partition hundreds of expressed proteins into six sectors with shared metabolic function and physiological response^4^. To accommodate the increased growth rates in the evolved strains, expression of metabolic enzymes undergoes sector-specific adaptation to enable increased fluxes. We find that catabolic proteins adapt by increasing the total enzyme abundance, whereas anabolic and glycolytic proteins exhibit decreased free-enzyme pools. We propose that flux-dependent regulation^7^ and substrate saturation^8^ can account for the sector-specific remodeling.

---

Laboratory evolution experiments provide a controlled environment within which adaptation can be tracked with single-nucleotide precision^1,6^. Despite the depth of data, there is no consensus framework for predicting the trajectory of adaptation, nor de-convolving how the genomic changes confer increased fitness. Part of the challenge comes from the large number of mutations that accumulate over the course of the experiment, and the possible interdependence of each of these mutations on all others in the evolutionary history of the organism^9,10^. Yet the recurrence of a common collection of mutations in parallel lineages hints at a set of simplifying principles^11,12^.

Physiological constraints can shape gene expression^5^. In *Escherichia coli*, one primary physiological constraint is the near-constant total protein concentration, reflecting the high metabolic cost of biosynthesis of the proteome. As a consequence, if one protein increases in concentration, then the concentrations of other proteins must decrease to accommodate the change. In response to growth inhibition, the concentration (or, equivalently, the protein mass fraction) of groups of proteins exhibit concerted positive or negative correlation with growth rate serving to partition the proteome into coarse-grained ‘sectors’ that share common functions^4,13^. Taken together, the constraint on total protein concentration and the coarse partitioning of the proteome define a set of physiological constraints that shape the response of the organism to metabolic challenges^5,13,14^.

For example, in response to translation-inhibition, ribosome and ribosome-affiliated proteins increase in mass fraction, with a concomitant decrease in the mass fraction of most other proteins. This set of co-regulated, translation-associated proteins defines what is referred to as the R-protein sector ^3–5^. Similarly, metabolic limitations (including catabolic and anabolic limitations) serve to define other protein groups responding in concert, resulting in a partitioning of the proteome comprised of six sectors^4,13^. The exponential growth rate can be decomposed as a flux-balance among these sectors, effectively treating each sector as a single enzyme with a lumped catalytic constant that quantifies how strongly changes in the sector protein abundance affects the growth rate^4^.

Proteome partitioning constraints have been used to predict metabolic response on laboratory timescales^5,14,15^, and on short evolutionary timescales to predict the spectrum of antibiotic-resistant mutants^16^. Here, we use quantitative proteomics applied to Lenski’s laboratory evolved strains to determine how adaptation to growth in glucose over the course of 40 000 generations is reflected in the proteome partitioning of the bacterium. There is a monotonic increase in the exponential growth rate of the bacteria with generation, necessitating a monotonic increase in the metabolic flux over the course of adaptation. We find that this increase in flux is accommodated differently in different sectors: the catabolic sector increases its total protein fraction, whereas the glycolytic and anabolic sectors keep a constant proteome fraction that carries more flux by increasing substrate saturation. This is effectively a decrease in the amount of enzyme not carrying flux, primarily in enzymes responsible for glycolysis and amino acid synthesis. Interestingly, these enzymes are adjacent in the metabolic network to two major deletions that appear early in the adaptation, pyruvate kinase (*pykF*) and glutamate synthase (GOGAT, *gltBD*). In response to the deletions, it appears that the bacterium is modulating substrate saturation rather than enzyme abundance to increase metabolic flux, thereby freeing up precious space in a constrained proteome.

## Results

We have used strains isolated from the Ara-1 lineage of Lenski’s long-term evolution experiment^6^. Lenski’s adaptation protocol is to grow *E. coli* in a minimal medium containing glucose and citrate (which is not metabolized by any of the strains in this lineage) as the sole carbon sources. Every 24 hours, the cells are diluted 1:100 into fresh media. The relative fitness (assessed via competition against the ancestral strain over one growth cycle) and the doubling rate of the evolved strains increase monotonically with generation number (Fig. S1). After 20k generations, the genome carries 29 single-nucleotide polymorphisms (SNPs) and 16 deletion-insertion polymorphisms (DIPs). Subsequently, this line develops a ‘hypermutator’ phenotype with dramatically-elevated mutation rate so that by 40k generations the genome carries 627 SNPs and 26 DIPs^6^.

### Adaptation of the ribosome abundance

The RNA/protein ratio serves as a proxy for the ribosome concentration^3^. Using various nutrient combinations, but no perturbation to the protein translation rate, the RNA/protein ratio (and consequently the ribosome mass fraction) exhibits a linear positive correlation with the doubling rate (Fig. 1A; grey line). The growth-rate dependence in the abundance of charged ternary complex produces a strong growth-rate dependence on the peptide elongation rate^17^, and yet the result is a simple linear relationship that is mathematically-equivalent to partitioning the ribosome abundance into a flux-carrying fraction, **Δr**, translating at maximum rate^3^, and a growth-rate independent fraction, **r**_**min**_, that is proportional to the Michaelis constant associated with ternary-complex binding^18^. Under the adaptation conditions of the Lenski experiment, the maximum translation rate (proportional to the inverse of the slope) and ternary-complex binding affinity (proportional to the intercept) exhibit no significant change, even after 40k generations (Fig. 1A; grey stars).

**Figure 1:**
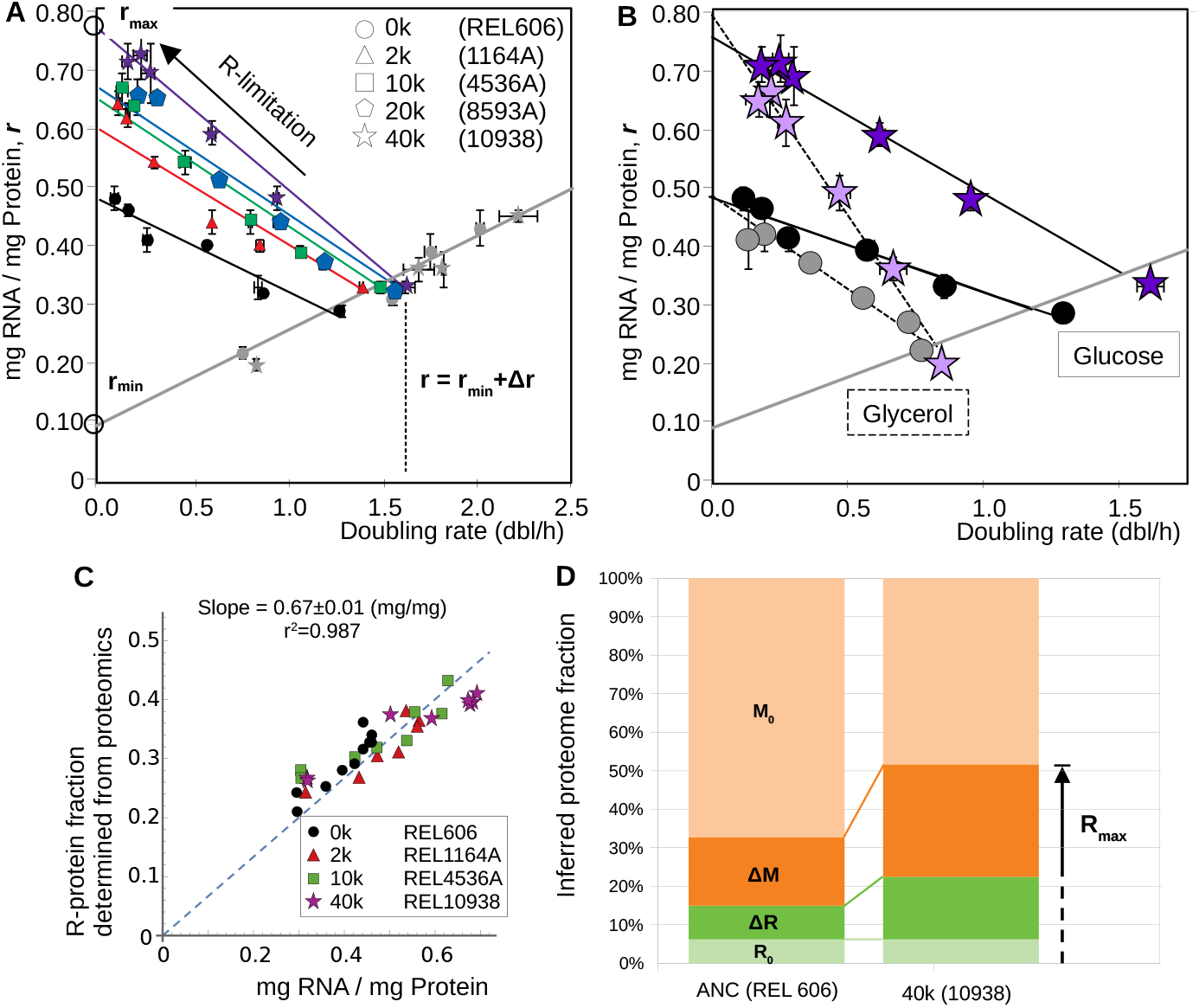
Effect of adaptation on ribosome abundance. **A**. The RNA/protein ratio is a proxy for the ribosome abundance. Over the course of adaptation, there is a decrease in the slope and a considerable increase in the maximum intercept under translation inhibition (R-limitation). Symbols denote the number of generation of adaptation; colored symbols denotes glucose minimal media with increasing concentrations of chloramphenicol, gray symbols are strains grown in various nutrient conditions (Table S1). Solid lines are the linear best-fit to the data and error bars denote one standard deviation over at least two biological replicates (Table S3). **B**. The increase in R-limited intercept is a generic feature that does not depend upon the carbon source. Here, the black and purple symbols are as in panel A; the gray and lavender correspond to R-limited growth in glycerol minimal media (Table S2). **C**. The RNA/protein ratio ***r*** is proportional the ribosome-affiliated protein fraction **R**, where **R**=0.67 x **r. D**. The nutrient-limited offset ***r*min** is attributed to incomplete saturation of the elongating ribosome with charged ternary complex^18^, effectively partitioning the ribosome pool into a non-flux carrying fraction **R0**=0.67 x ***rmin*** and a flux-carrying fraction **ΔR**=0.67 x (***r***-***r*min**). Upon translation-inhibition, the flux-carrying fraction expands, compressing the growth-dependent fraction of metabolic proteins **ΔM**. The nominal abundance of these metabolic proteins can be inferred from **ΔM**=0.67 x (***r*max**-***r***). Using this two-sector partition of the proteome, the primary effect of the adaptation is a decrease in the growth-rate independent fraction **M0**=1-**R**-**ΔM** allowing both growth-rate dependent sectors (**ΔR** and **ΔM**) space to increase.

When the bacteria are treated with the ribosome-targeting antibiotic chloramphenicol, the protein synthesis flux is decreased as ribosomes are inactivated^17^. For a fixed nutrient condition, the RNA/protein ratio exhibits a linear negative correlation with doubling rate, as the rate of ribosome synthesis is increased to counteract the inhibition (Fig. 1A; colored lines). The maximum intercept, **r**_**max**_, increases monotonically with generation number (Fig. 1A, colored lines). Fig. 1B shows that the intercept is identical whether the cells are grown in glucose (solid lines) or glycerol (dashed lines) minimal medium, suggesting that the intercept is not specific to growth in glucose.

The RNA/protein ratio data was corroborated by measuring the R-protein fraction using mass spectrometry. Proteomic mass spectrometry can quantify small changes in protein abundance across hundreds of proteins^4^. Samples were collected during exponential growth of the ancestral (REL606) and 40k strain (10938) in glucose minimal medium under nominal growth and translation-limitation (R-limitation). These were mixed with a heavy-nitrogen labeled standard grown in glucose minimal medium using ^15^NH_4_Cl as the sole metabolizable nitrogen source. Peptide counts, normalized relative to the standard, are used to estimate the protein mass fraction of a given protein^4^. As shown in Fig. 1C, the RNA/protein ratio is proportional to the R-protein fraction across the adaptation, with a proportionality constant of 0.67 x RNA/protein = R-protein fraction.

The R-proteins occupy a large fraction of the proteome. When they are upregulated in response to translation-inhibition (‘R-limitation’), the proteome partitioning constraint implies that the non-ribosomal protein fraction must decrease to accommodate the change^3,5^. The empirical proportionality between the RNA/protein ratio **r** and the R-protein fraction allows the RNA/protein data to be expressed in terms of protein mass fraction, providing a coarse-grained overview of how the adaptation affects the proteome partitioning.

The total R-protein fraction is composed of an offset **R**_**0**_ = 0.67 x **r**_**min**_ (that from Fig. 1A is approximately constant over the course of adaptation), and a growth-rate dependent fraction **ΔR**=0.67 x (**r**-**r**_**min**_). The remainder of the proteome, largely occupied by metabolic proteins^19^. In what follows, it will be useful to partition the metabolic protein fraction into two parts: one part that responds to growth rate changes and decreases under R-limitation which we call **ΔM**, and a growth-rate independent remainder **M**_**0**_. Under R-limitation, the R-protein fraction increases (with a subsequent, and complementary, decrease in the growth-rate-dependent non-ribosomal fraction **ΔM**) until it reaches a maximum allowable fraction **R**_**max**_ = 0.67 x **r**_**max**_. From the nominal R-protein fraction and the R-limitation intercept, we can infer both the growth-dependent metabolic protein fraction^3^ **ΔM**=0.67 x (**r**_**max**_-**r**), and the remainder **M**_**0**_=1-**R**_**0**_-**ΔR**-**ΔM**.

Fig. 1D illustrates the effect of adaptation on this coarse partitioning of the proteome. The ancestral strain (REL606) has a comparatively small intercept **r**_**max**_ (and correspondingly large fraction **M**_**0**_). After 40k generations of glucose-adapted growth, the intercept **r**_**max**_ is larger due to a decrease in the offsets **M**_**0**_. The growth-rate dependent ribosomal and metabolic fractions (**ΔR** and **ΔM**) both increase over 40k generations.

### Adaptation of proteome sectors

Previous work established a coarse-grained partitioning of the proteome based upon response to growth limitations^4^. The partitioning consists of six sectors that exhibit increased protein fraction under: translation limitation (R), catabolic limitation (C), anabolic limitation (A), both catabolic and anabolic limitation (S), a growth-rate independent sector (O), and the growth-rate-dependent, but limitation independent, sector (U). In addition to shared physiological response, proteins in each sector share common metabolic roles (Fig. 2A).

**Figure 2:**
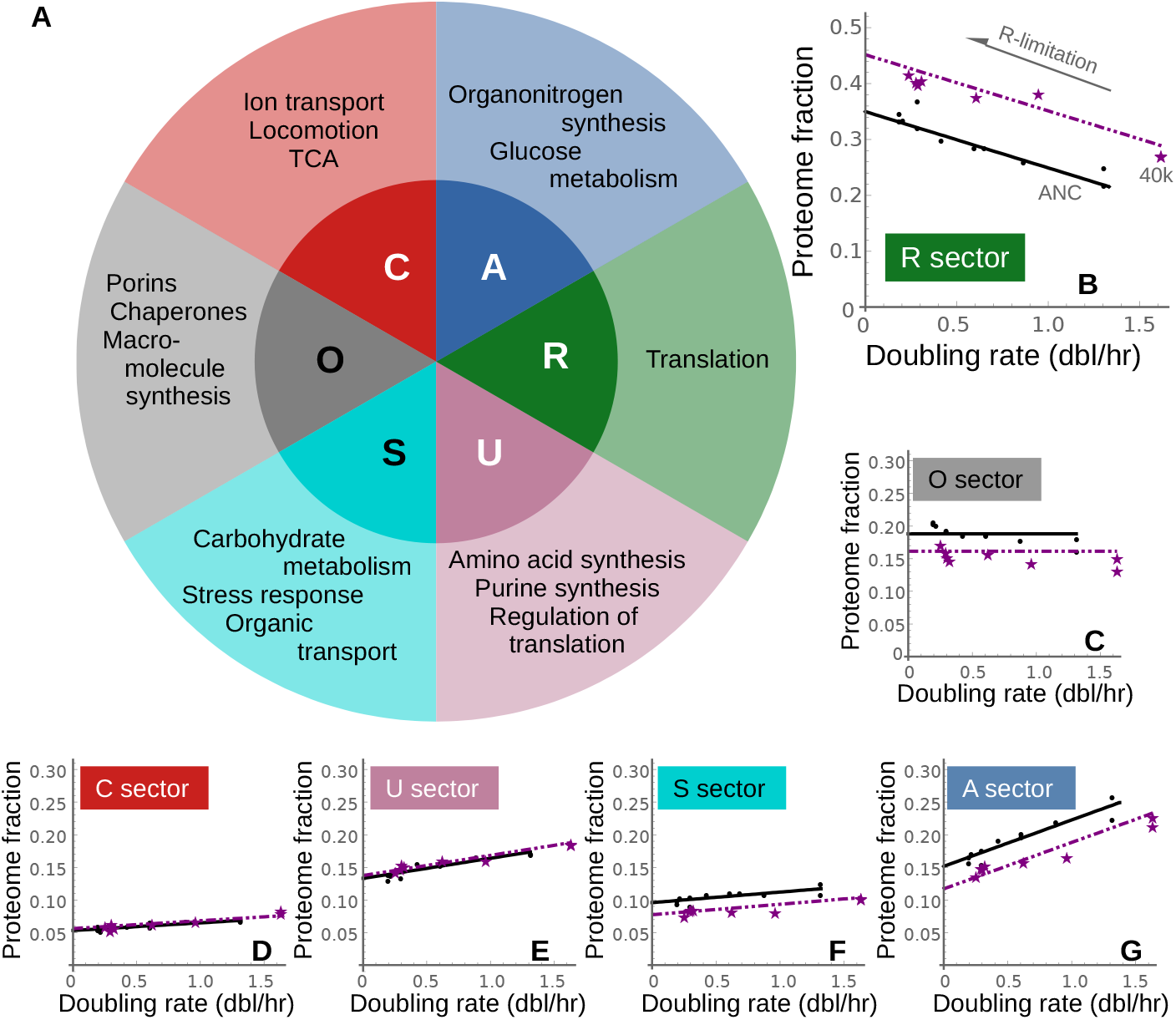
Effect of adaptation on proteome partitioning. **A**. Partitioning of the proteome based upon physiological response to growth inhibition results in six sectors, with shared function among proteins in each sector^4^. **B**. After 40k generations of adaptation, the R-sector exhibits the same increase in the R-limited intercept as the RNA/protein ratio shown in Fig 1A. **C**. The growth-rate independent O-sector exhibits decreased expression levels. (**D**,**E**) There is little change in the R-limited response of the C and U sectors. (**F**,**G**) The S- and A-sectors appear to adapt via changes in intercept without changes in slope. The black lines are best-fit to the ancestral data; the purple dashed lines are the best-fit across the 40k strain assuming no change in slope (such that the intercepts across all six sectors sum to 1, Table S4).

The R-protein fraction behaves similarly to the RNA/protein ratio, although there is no appreciable change in the slope of the R-limitation line between the ancestral and 40k strains (Fig 2B). Adaptation affects the R-limited behavior of the non-ribosomal proteome sectors in different ways. The growth-rate-independent O-sector exhibits adaptation by decreasing the expression level (Fig. 2C). The C- and U-sectors lie upon the same R-limitation line in both the ancestral and 40k strains (Fig. 2D and 2E). The remaining A- and S-sectors exhibit adaptation through change in intercept alone, without any apparent change in the slope (Fig. 2F and 2G). In Figs. 2B-G, the linear fits to the 40k data (purple) are done by adjusting the intercept subject to the constraint that the R-limited intercepts across all six sectors sum to one (Supplemental Table S4). The combined behavior of the non-ribosomal protein sectors is consistent with the coarse partitioning derived from the RNA/protein ratio insofar as the growth-rate dependent fraction of each sector increases after adaptation (Fig. S2).

Adaptation at the level of individual proteins is summarized in Fig. 3A, illustrating the correlation between changes in nominal protein expression (at a growth rate of 1.3 dbl/h) and changes in the R-limited intercept between the ancestral and 40k strains. Proteins for each sector are shown as individual dots, while ellipses capture 68% of the data. Most data lie on the diagonal, implying that most proteins adapt by changing their intercepts, rather than their slopes. The solid circles, representing for each sector the average changes across proteins irrespective of their abundance, match well the predicted changes obtained from the proteome-sector fits shown in Figs. 2B-G.

**Figure 3:**
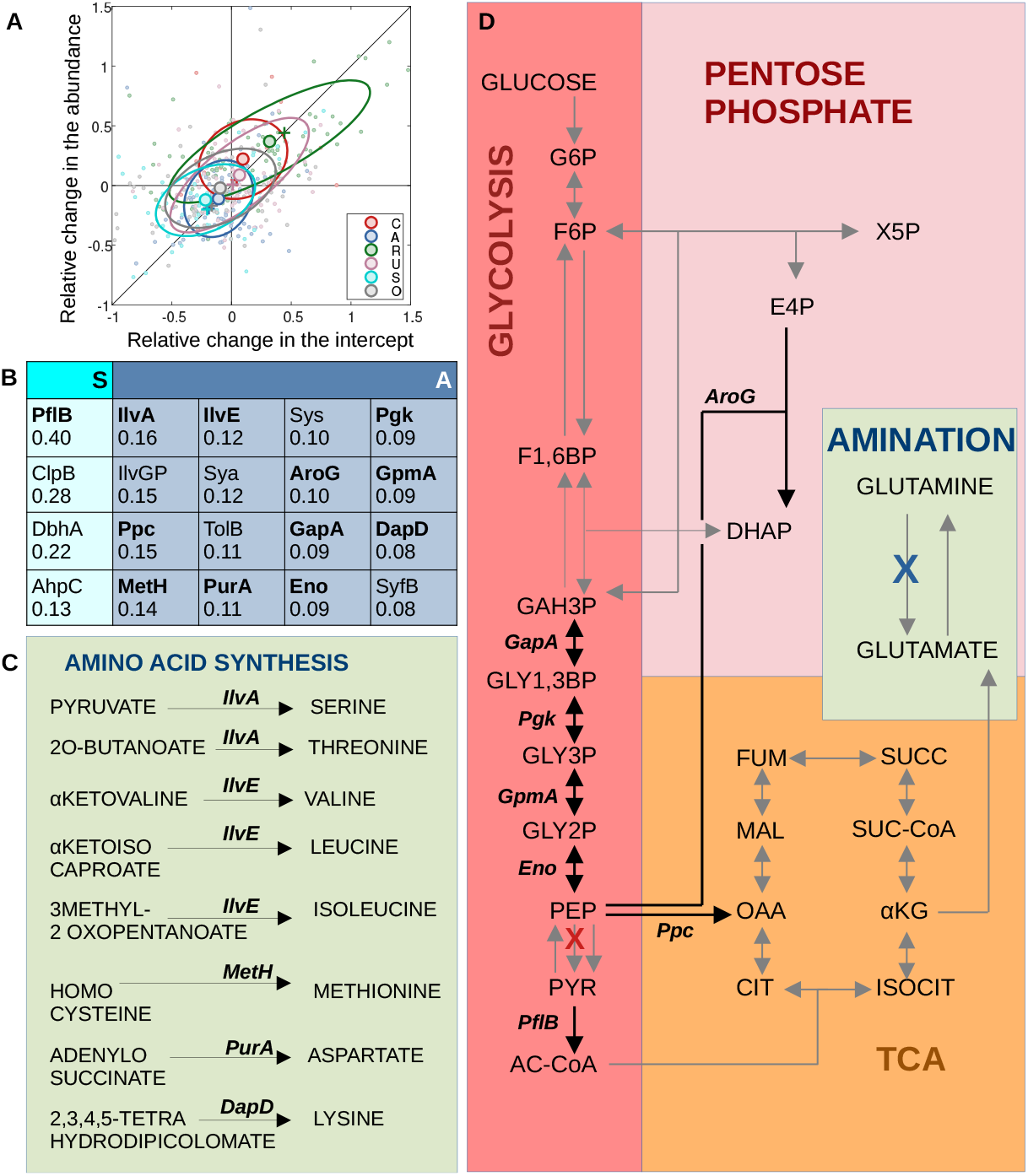
Protein expression changes close to deletions of PykF and GltBD. **A**. Sector behavior is largely recapitulated at the level of individual proteins. The relative change in the nominal protein fraction (normalized to growth at 1.3 dbl/h) is compared to the relative change in the intercept for individual proteins. Those points lying on the diagonal exhibit no change in slope upon adaptation. Large circles correspond to the average over each sector, along with one-standard-deviation ellipses; these can be compared to the changes (crosses) predicted by the sector fits shown in Fig. 2. Low abundance proteins (less than 400 ppm) were not included in the figure and represent less than 15% of the detected protein fraction. **B**. Minimal set of genes that explain 50% of the observed reduction in the growth-independent fraction of the S- and A-sector proteins. The numbers correspond to the absolute change in R-limited intercept protein mass fraction between the ancestral and 40k strains (expressed as percent). Changes in the R-limited intercepts of enzymes involved in (**C**) amino acid synthesis and (**D**) carbon metabolism cluster around two deletions occurring early in adaptation: pyruvate kinase (*pykF*, red cross) and GOGAT (*gltBD*, blue cross). Enzymes in bold correspond with those shown panel B. Individual protein plots appear in Fig. S3. The metabolic map was redrawn from the KEGG database^38^.

Finally, we investigated which proteins had the largest contribution to the adaptation observed at the protein sector level. The decrease in the O-sector intercept is primarily due to a regulated decrease in the abundance of the porin OmpF^20^. Highly expressed proteins belonging to the A- and S-sector also adapted primarily via changes in intercept. Fig. S3 shows the growth rate dependence for the proteins accounting for most of the observed change in the sector-level intercept; their contribution to the total change of the sector intercept is summarized in Fig. 3B. The changes in the S-sector proteins are confined to a short list, including the glycolytic enzyme PflB. By contrast, the A-sector proteins comprise small changes distributed among many genes. The S- and A-sector proteins responsible for the majority of the proteome change are clustered in the metabolic map near two highly-abundant, high-flux-carrying proteins, both of which were deleted early in the adaptation: glutamate synthase (GOGAT *gltBD*; blue cross, Fig. 3D) and pyruvate kinase (*pykF*; red cross). The bold arrows and enzymes in Figs. 3C and 3D correspond to the bold entries in Fig. 3B.

## Discussion

The R-limited behavior of the proteome sectors can be rationalized by considering the behavior of an irreversible enzyme. For a simple enzymatic reaction, with substrate in excess of the enzyme (that is, the ‘enzyme-limited regime’), the rate of product formation is given by the familiar Michaelis-Menten form (Fig. 4A). There is a dynamic equilibrium between the substrate-bound active complex **E**_**a**_ and the free enzyme **E**_**f**_, but the flux is strictly proportional to the active complex concentration (Fig. 4B).

**Figure 4:**
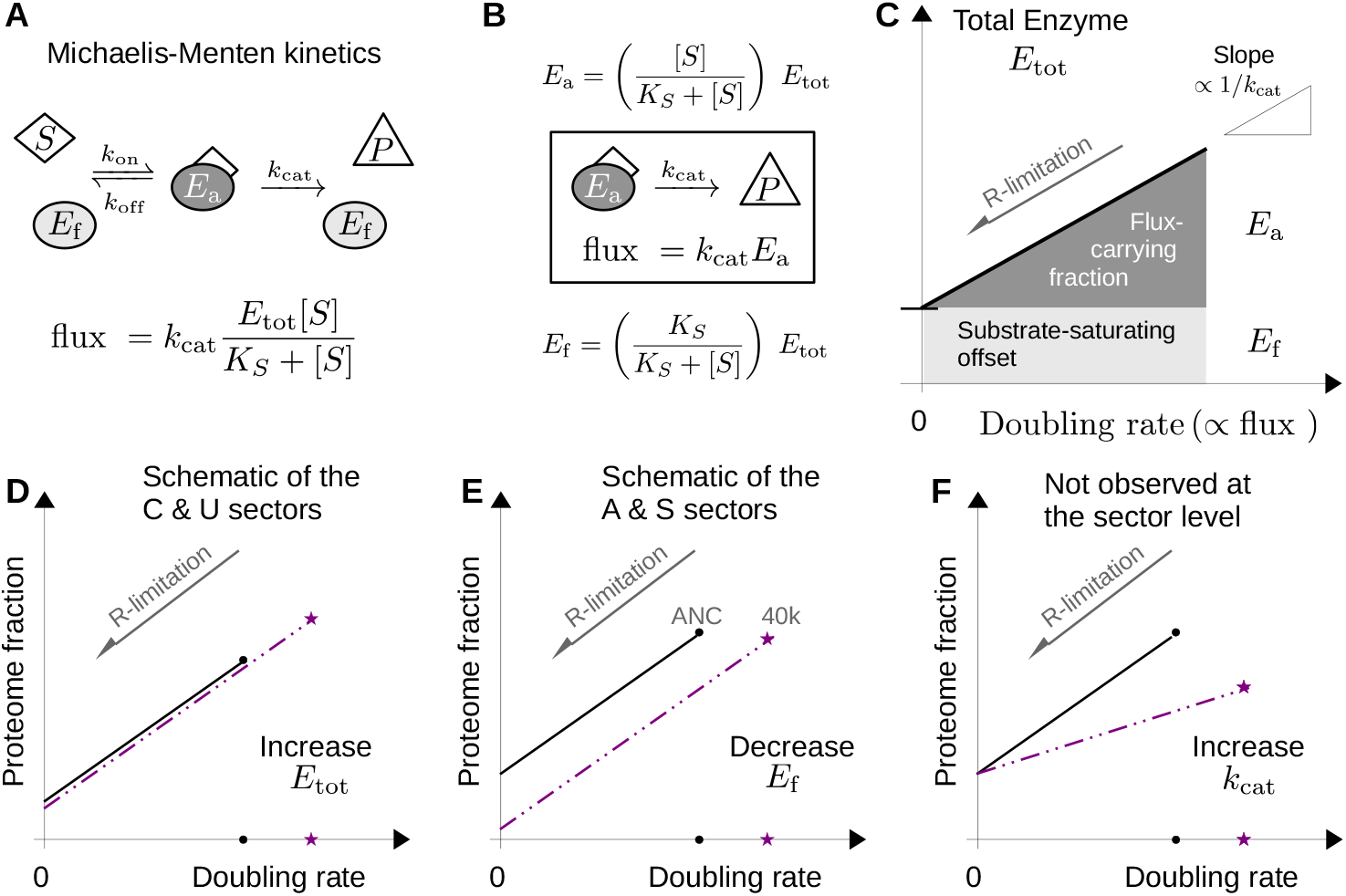
Linearity in the proteome fractions. **A**. For an irreversible enzymatic reaction, the flux is proportional to the concentration of enzyme engaged in active complex with the substrate. If the substrate concentration **[S]** far exceeds the total concentration of enzyme **Etot**=**Ea**+**Ef**, then the active complex concentration **Ea** takes the familiar Michaelis-Menten form^38^. **B**. Distinguishing between the active complex **Ea** and the free enzyme **Ef** provides a rationalization for the proteomic data. **C**. For many enzymatic reactions, flux is proportional to the doubling rate^23^. Inhibiting translation (R-limitation) attenuates the doubling rate without affecting the catalytic constant kcat^3,4^. Consequently, the R-limited response of an individual enzyme provides an estimate of the active flux-carrying enzyme abundance **Ea**, the free enzyme abundance **Ef** (intercept) and the catalytic constant **kcat** (reciprocal slope). **D**. Adaptation in the C and U-sectors is analogous to an increase in the total enzyme abundance **Etot. E**. In contrast, adaptation in the A- and S-sectors is analogous to a decrease in the free enzyme **Ef. F**. At the coarse-grained level, we did not observe any change in the slope of the R-limited response, analogous to an increased catalytic efficiency **kcat**.

Previous work has quantified subtle changes in nominal protein expression in the adapted strains by using 2D gels^21^ or ribosome density^22^. What is different here is that by coupling proteomics with R-limitation, we are able to estimate how much of the nominal protein is flux-carrying. For many reactions, the flux is proportional to the doubling rate^23^: cells growing twice as quickly must convert metabolic intermediates twice as quickly. The utility of R-limitation is that it provides a method for modulating the doubling rate (and therefore the reaction fluxes) by modulating the protein synthesis rate without affecting the catalytic constants of the metabolic enzymes. Consequently, as the doubling rate is decreased via translation-inhibition, the abundance of individual enzymes decreases in linear proportion by directly attenuating the abundance of active (flux-carrying) enzyme **E**_**a**_ (Fig. 4C). The intercept of the R-limited response corresponds to the free-enzyme abundance **E**_**f**_, and the slope is proportional to the reciprocal of the catalytic constant **k**_**cat**_. A similar analysis can be applied to reversible enzymes (Fig. S4).

The R-limited response of a given proteome sector is a weighted average of the enzymes that make up that sector^24^. After 40k generations, the doubling rate is increased necessitating increased flux through each sector; three possible scenarios are shown schematically in the lower panels of Fig. 4. For the catabolic C- and U-sector proteins, the R-limited behavior is consistent with an increase in flux mediated primarily by increasing the total protein abundance (Fig. 4D). If the intercept and slope remain unchanged, an increase in the total enzyme abundance directly increases the flux-carrying fraction. The R-limited response shown in Fig. 4D can be achieved through feed-forward flux-controlled regulation^5,7^; either at the level of individual genes or via a master regulator^25^.

By contrast, the biosynthetic and glycolytic S- and A-sector proteins exhibit R-limited behavior that is consistent with a decrease in the intercept without an apparent change in the nominal enzyme abundance or the slope (Fig. 4E). For a single enzyme, this scenario would correspond to a decrease in the free enzyme abundance. It is less clear how to achieve the observed growth-rate dependence, although for a single enzyme, the ratio of the active and free enzyme abundance is equal to the scaled substrate concentration **[S]**/**K**_**s**_ where **K**_**s**_ is the Michaelis constant for the reaction (Fig. S4). The A- and S-sectors behave as though the substrate saturation is increased under nominal growth conditions in the 40k strain. Based upon the topology of the metabolic network (Fig. 3D), it appears that the PykF/GlyBD deletions create bottlenecks that increase substrate saturation, allowing for an increase in catalyzed flux without the need for increased protein expression. Flux regulation via substrate saturation is observed under conditions of anabolic limitation (for example by titrating the abundance of the GltBD enzyme, GOGAT^26^).

Strong selective pressure directed toward increasing the flux through a single reaction can result in mutations that improve enzyme performance^27,28^, which would appear as a reduced slope when plotting the enzyme levels against the growth rate (Fig. 4F), yet this was not observed in any of the coarse-grained sectors. For the R-sector, the lack of change in the slope in nutrient-limiting conditions suggests that the ribosome elongation rate is unchanged after adaptation.

For growth in glucose, the PykF/GltBD deletions appear to be best rationalized by the proteome remodeling that occurs in their absence, although it is less clear how this remodeling affects growth on other substrates^29^. The diminished function (or occasionally deletion) of pyruvate kinase (*pykF*) is commonly observed in the adaptation of *E. coli* to growth in glucose minimal media^6,10,12^, and in the directed evolution of isobutyrate^30^ and L-serine^31^ producing strains. Consistent with our observed decrease in the non-flux carrying intercepts of the A- and S-sector proteins, deletion of *pykF* does increase glycolytic substrates (notably glucose-6-phosphate and fructose-6-phosphate), as well as reducing cellular pyruvate levels^32^. Because pyruvate is one of the primary reporters of carbon flux in *E. coli*^14,33^, and we expect the loss of *pykF* in the Lenski strains to result in impaired carbon catabolite repression.

The R-limited proteomic response is a tool with application outside of the coarse-grained physiological characterization we have done. Applied to individual enzymes, the interpretation outlined in the upper panels of Fig. 4 provide a framework for estimating *in situ* enzymatic activity. Taking the ratio of the flux to the total enzyme concentration, the enzyme activity is **k**_**cat**_/(1+**E**_**f**_/**E**_**a**_) where all three parameters are fully-determined by the R-limited response. At present, the *in situ* activity of individual metabolic enzymes is estimated using isotope-tracing for the flux and quantitative protein abundance; the R-limited behavior of individual proteins provides a convenient complementary methodology.

Framing adaptation dynamics in terms of a coarse-grained physiological model can be applied more generally to other microorganisms, including engineered ‘minimal genome’ strains^34^. In the Lenski-adapted strains, the R-limited intercepts exhibit a relative change with generation number that is about 100x more rapid than changes in the doubling rate. As a consequence, significant remodeling of the proteome can be observed on a timescale of several weeks of adaptation, rather than several decades, allowing for comparatively-convenient exploration of a variety of laboratory evolution scenarios. Furthermore, systematic comparison of R-limited proteomics across bacterial species would provide a complementary tool to explore how expression is shaped by the shared evolutionary history of the species, rather than adaptation to specific ecological niches on (comparatively) shorter timescales. The gram-positive bacterium *B. subtilis* and the gram-negative *V. natriegens* were recently found to express a majority of metabolic proteins to similar levels despite their distinct evolutionary histories^35^, but how much of the protein is flux-carrying remains unknown.

Our focus has been on the physiological consequences of adaptation manifest in exponential growth. The Lenski strains spend about seven hours a day in that state, the other eighteen hours are spent transitioning in and out of stationary phase, and consuming waste acetate. A complete picture of physiological adaptation to this adaptation regimen would necessarily include these other growth states. Nevertheless, the observed changes in proteome partitioning can be largely understood in terms of adjusting metabolic flux through changes in protein expression following deletion events. Further, these proteomic changes could not be deduced from the available genome sequences during adaptation, highlighting the importance of the expressed proteome as an energetically-costly carrier of metabolic flux.

## Methods

### Strains and growth conditions

We have used Lenski’s Ara-1 lineage derived from a B strain ancestor (REL 606), and isolated after 2K (strain REL1164A), 10K (REL4536A), 20K (REL8593A) and 40K (REL10938) generations of growth as described previously^6^. In this study, strains were grown in MOPS minimal medium^36^ (Teknova, M2106) supplemented with 0.2% (w/v) glucose or 0.2% (v/v) glycerol as the carbon source. For nutrient-modulated growth (Fig. 1A) the minimal medium was enriched using 0.2% (w/v) casamino acids (Fisher), or nucleotides and amino acids (Teknova,M2103 and M2104) as indicated in Table S1 (denoted by CAA [casamino acids] and RDM [rich defined medium], respectively). Translation-limited (R-limited) growth was achieved by adding chloramphenicol to the growth medium at sublethal concentrations (0-12 μM) as indicated in Tables S1 and S2. Strains were grown in test tubes at 37°C in a water-bath shaker (Thermo-Fisher MaxQ7000), agitated at 250 r.p.m. for aeration. Doubling rate was determined by following the rate-of-change in scattered light at 600 nm (OD600) using a Thermo-Fisher BioMate3S UV-visible spectrophotometer.

### RNA and protein extraction assays

The total RNA (μg/mL/OD600) was determined by precipitation in perchloric acid^37^, with modifications as previously described^14^. The total protein (μg/mL/OD600) was determined using a Lowry kit (Sigma-Aldrich, TP0300), and bovine serum albumin (BSA) as the standard.

### Quantitative proteomics

Samples were collected and processed as previously described^4^. The heavy-nitrogen standard was prepared by combining 1:1 (w/w) protein isolated from the ancestral strain grown in glucose minimal medium (20 mM ^15^NH_4_Cl) with protein isolated from the 40k strain grown in glucose minimal medium under R-limitation (20 mM ^15^NH_4_Cl and 12 μM chloramphenicol).

### Data availability

Raw mass spectral data is deposited to massIVE, with the accession code MSV000087313 (ftp://MSV000087313@massive.ucsd.edu) or available through the Proteomexchange (http://www.proteomexchange.org/) via the accession code PXD025666. These will be made publicly-accessible upon publication. All other data are available in the supplementary tables.

## Supporting information

Peptide counts

## Acknowledgements

We are grateful to Richard Lenski for providing the glucose-adapted strains. This work is supported by the NIH through grant R35GM136412 (JRW) and by NSERC through grant 2016-03658 (MS).

## Author contributions

Conceptualization: M.S. Investigation: M.S. (growth, RNA and protein), M.S. and M.M. (data analysis), V.P and J.R.W. (quantitative proteomics). Writing - original draft: M.S. Writing - review and editing: M.S., J.R.W. and M.M. All authors reviewed the results and approved the final version of the manuscript.

## Display captions

**Figure S1**: The relative fitness and the doubling rate monotonically increase with generation number. Fitness data is from Barrick *et al*.^6^

**Figure S2**: The proteome allocation in glucose minimal medium for the ancestral (REL606) and 40k strain (10938) using the more detailed taxonomy proposed by Hui et al.^4^ (described in Fig 2A of the main text) recapitulates the allocation inferred from the RNA/protein measurements shown in Fig. 1C of the main text. The growth-dependent fractions of the proteome increases from 30% of the proteome in the ancestral strain to about 40% in the adapted strain, accommodated primarily by a decrease in the growth-rate independent fraction of the A sector and a decrease in the non-flux carrying O sector. The bars on the far-right correspond to the taxonomy in Fig. 1C, where the protein fractions are inferred from the RNA/protein ratio. As in Fig. 1C, the growth-rate dependent fractions of ribosomal (**ΔR**, green) and metabolic (**ΔM**, orange) proteins increase concomitantly with the increased growth rate over the course of adaptation, at the expense of a diminished growth-rate independent fraction of metabolic proteins (**M**_**0**_, pale orange). Sectors with similar behaviour (C/U and A/S) have been lumped together to improve the readability of the figure.

**Figure S3**: Proteome fractions of individual S- and A-sector proteins under R-limitation in the ancestral strain (REL606, black) and the 40k strain (10938, purple) as a function of the exponential growth rate. The proteins correspond to the entries of Fig. 3B in the main text.

**Figure S4**: Summary of Klumpp *et al*.^18^ and Dourado *et al*.^8^. **A)** Assuming reversible Michaelis-Menten kinetics, the flux through the reaction is proportional to a fraction of the total protein concentration, **E**_**tot**_. For many metabolic reactions, the flux is proportional to the growth rate, and so the flux-carrying protein fraction **E**_**a**_ is assumed to correspond with the growth-rate dependent protein fraction observed in the proteomic data. Consequently, the growth-independent offset is given by the free enzyme fraction **E**_**f**_. The ratio of the two fractions, **E**_**a**_/**E**_**f**_, is equal to the substrate saturation **[S]**/**K**_**s**_, where **K**_**s**_ is the Michaelis constant for the reaction. **B)** Though the expression is more complicated for a reversible enzyme, the non-flux carrying fraction **E**_**f**_ can again be reduced by increasing the substrate saturation **[S]**/**K**_**s**_.

## References

1. Sandberg, T. E., Salazar, M. J., Weng, L. L., Palsson, B. O. & Feist, A. M. The emergence of adaptive laboratory evolution as an efficient tool for biological discovery and industrial biotechnology. Metab. Eng. 56, 1–16 (2019).

2. Lenski, R. E. Convergence and Divergence in a Long-Term Experiment with Bacteria. Am. Nat. 190, S57–S68 (2017).

3. Scott, M., Gunderson, C. W., Mateescu, E. M., Zhang, Z. & Hwa, T. Interdependence of cell growth and gene expression: origins and consequences. Science 330, 1099–1102 (2010).

4. Hui, S. et al. Quantitative proteomic analysis reveals a simple strategy of global resource allocation in bacteria. Mol. Syst. Biol. 11, 784 (2015).

5. Scott, M. & Hwa, T. Shaping bacterial gene expression by physiological and proteome allocation constraints. Nat. Rev. Microbiol. 21, 327–342 (2023).

6. Barrick, J. E. et al. Genome evolution and adaptation in a long-term experiment with Escherichia coli. Nature 461, 1243–1247 (2009).

7. Litsios, A., Ortega, Á. D., Wit, E. C. & Heinemann, M. Metabolic-flux dependent regulation of microbial physiology. Curr. Opin. Microbiol. 42, 71–78 (2018).

8. Dourado, H., Mori, M., Hwa, T. & Lercher, M. J. On the optimality of the enzyme–substrate relationship in bacteria. PLOS Biol. 19, e3001416 (2021).

9. Khan, A. I., Dinh, D. M., Schneider, D., Lenski, R. E. & Cooper, T. F. Negative Epistasis Between Beneficial Mutations in an Evolving Bacterial Population. Science 332, 1193–1196 (2011).

10. Peng, F. et al. Effects of Beneficial Mutations in pykF Gene Vary over Time and across Replicate Populations in a Long-Term Experiment with Bacteria. Mol. Biol. Evol. 35, 202–210 (2018).

11. Wang, X., Zorraquino, V., Kim, M., Tsoukalas, A., & Tagkopoulos, I. Predicting the evolution of Escherichia coli by a data-driven approach. Nat. Commun. 9, 3562 (2018).

12. Phaneuf, P. V., Gosting, D., Palsson, B. O. & Feist, A. M. ALEdb 1.0: a database of mutations from adaptive laboratory evolution experimentation. Nucleic Acids Res. 47, D1164–D1171 (2019).

13. Mori, M. et al. From coarse to fine: the absolute Escherichia coli proteome under diverse growth conditions. Mol. Syst. Biol. 17, e9536 (2021).

14. You, C. et al. Coordination of bacterial proteome with metabolism by cyclic AMP signalling. Nature 500, 301–306 (2013).

15. Erickson, D. W. et al. A global resource allocation strategy governs growth transition kinetics of Escherichia coli. Nature 551, 119–123 (2017).

16. Pinheiro, F., Warsi, O., Andersson, D. I. & Lässig, M. Metabolic fitness landscapes predict the evolution of antibiotic resistance. Nat. Ecol. Evol. 5, 677–687 (2021).

17. Dai, X. et al. Reduction of translating ribosomes enables Escherichia coli to maintain elongation rates during slow growth. Nat. Microbiol. 2, 1–9 (2016).

18. Klumpp, S., Scott, M., Pedersen, S. & Hwa, T. Molecular crowding limits translation and cell growth. Proc. Natl. Acad. Sci. U. S. A. 110, 16754–16759 (2013).

19. Mori, M., Cheng, C., Taylor, B. R., Okano, H. & Hwa, T. Functional decomposition of metabolism allows a system-level quantification of fluxes and protein allocation towards specific metabolic functions. Nat. Commun. 14, 4161 (2023).

20. Crozat, E. et al. Altered Regulation of the OmpF Porin by Fis in Escherichia coli during an Evolution Experiment and between B and K-12 Strains. J. Bacteriol. 193, 429–440 (2011).

21. Pelosi, L. et al. Parallel Changes in Global Protein Profiles During Long-Term Experimental Evolution in Escherichia coli. Genetics 173, 1851–1869 (2006).

22. Favate, J. S., Liang, S., Cope, A. L., Yadavalli, S. S. & Shah, P. The landscape of transcriptional and translational changes over 22 years of bacterial adaptation. eLife 11, e81979 (2022).

23. Varma, A. & Palsson, B. O. Stoichiometric flux balance models quantitatively predict growth and metabolic by-product secretion in wild-type Escherichia coli W3110. Appl. Environ. Microbiol. 60, 3724–3731 (1994).

24. Jun, S., Si, F., Pugatch, R. & Scott, M. Fundamental principles in bacterial physiology—history, recent progress, and the future with focus on cell size control: a review. Rep. Prog. Phys. 81, 056601 (2018).

25. Kochanowski, K. et al. Functioning of a metabolic flux sensor in Escherichia coli. Proc. Natl. Acad. Sci. 110, 1130–1135 (2013).

26. Kochanowski, K. et al. Global coordination of metabolic pathways in Escherichia coli by active and passive regulation. Mol. Syst. Biol. 17, e10064 (2021).

27. Quandt, E. M. et al. Fine-tuning citrate synthase flux potentiates and refines metabolic innovation in the Lenski evolution experiment. eLife 4, e09696 (2015).

28. Herring, C. D. et al. Comparative genome sequencing of Escherichia coli allows observation of bacterial evolution on a laboratory timescale. Nat. Genet. 38, 1406–1412 (2006).

29. Leiby, N. & Marx, C. J. Metabolic Erosion Primarily Through Mutation Accumulation, and Not Tradeoffs, Drives Limited Evolution of Substrate Specificity in Escherichia coli. PLoS Biol. 12, e1001789 (2014).

30. Lennen, R. M. et al. Laboratory evolution reveals general and specific tolerance mechanisms for commodity chemicals. Metab. Eng. 76, 179–192 (2023).

31. Mundhada, H. et al. Increased production of L-serine in Escherichia coli through Adaptive Laboratory Evolution. Metab. Eng. 39, 141–150 (2017).

32. Al Zaid Siddiquee, K., Arauzo-Bravo, M. J. & Shimizu, K. Metabolic flux analysis of pykF gene knockout Escherichia coli based on 13C-labeling experiments together with measurements of enzyme activities and intracellular metabolite concentrations. Appl. Microbiol. Biotechnol. 63, 407–417 (2004).

33. Okano, H., Hermsen, R., Kochanowski, K. & Hwa, T. Regulation underlying hierarchical and simultaneous utilization of carbon substrates by flux sensors in Escherichia coli. Nat. Microbiol. 5, 206–215 (2020).

34. Juhas, M., Reuß, D. R., Zhu, B. & Commichau, F. M. Bacillus subtilis and Escherichia coli essential genes and minimal cell factories after one decade of genome engineering. Microbiology 160, 2341–2351 (2014).

35. Lalanne, J.-B. et al. Evolutionary Convergence of Pathway-Specific Enzyme Expression Stoichiometry. Cell 173, 749–761.e38 (2018).

36. Neidhardt, F. C., Bloch, P. L. & Smith, D. F. Culture Medium for Enterobacteria. J. Bacteriol. 119, 736–747 (1974).

37. Benthin, S., Nielsen, J. & Villadsen, J. A simple and reliable method for the determination of cellular RNA content. Biotechnol. Tech. 5, 39–42 (1991).

38. Kanehisa, M., Sato, Y. & Kawashima, M. KEGG mapping tools for uncovering hidden features in biological data. Protein Sci. 31, 47–53 (2022).

39. Briggs, G. E. & Haldane, J. B. S. A Note on the Kinetics of Enzyme Action. Biochem. J. 19, 338–339 (1925).

